# Solvatochromic reporter to image plasma membrane order leaflet by leaflet reveals a highly asymmetric bilayer locally modulated by transbilayer interactions

**DOI:** 10.1101/2024.07.23.604763

**Authors:** Chandrima Patra, Muzamil Samad, Thomas van Zanten, Maninder Singh, Ram A Vishwakarma, Parvinder Pal Singh, Satyajit Mayor

## Abstract

The plasma membrane (PM) of cells is an asymmetric bilayer composed of a complex mixture of lipids and proteins. The emergent biophysical properties of its leaflets and their consequence on cellular function remain unexplored owing to the limitation in available probes. In this manuscript, we report on the design and characterisation of a PM-localised solvatochromic probe, C3L, based on the established reporter Laurdan. C3L is retained at the inner-leaflet due to flippase and scramblase activity, as determined by Bovine Serum Albumin-based back exchange experiments. By measuring the Generalized Polarization of C3L we are able to determine membrane order, leaflet-by-leaflet at the PM. Our results reveal that the PM is made of a tightly-packed outer/exofacial-leaflet in contrast to a disordered inner/cytoplasmic-leaflet. This differential packing is established by the maintenance of lipid asymmetry, the presence of specific lipids, Sphingomyelin in the outer-leaflet and Phosphatidylserine at the inner-leaflet, and exaggerated during mesenchymal to epithelial transition. We find spatial signatures of local modulation of the inner leaflet by protein scaffolds which promote trans-bilayer coupling of lipids. C3L thus informs on steady state lipid organisation at the asymmetric membrane while allowing further exploration of spatial regulation of lateral heterogeneity at each leaflet.

## INTRODUCTION

The plasma membrane (PM) is a ∼5nm thick asymmetric bilayer demarcating the cell from its external milieu, composed of proteins and lipids that are unequally distributed across the two leaflets^1^. This asymmetric lipid composition is established by the lipid-synthesis and remodelling programs, vesicular and non-vesicular transport and the enzymatic activity of ATP driven flippases, floppases and the ATP-independent scramblases at the membrane^2,3^. Hundreds of lipid species that consequently populate this membrane are distributed in a laterally heterogenous manner to form membrane domains^4^. These domains are formed by lateral interactions of lipids, augmented by the interaction of protein components with the juxtaposed cytoskeleton or a combination of both^5–8^. Current understanding suggests these domains can modify the output of signalling networks via the modulation of organization of receptors at the membrane thus allowing the cell to respond to intracellular and extracellular cues^9–11^. An understanding of the lipid environment and the resultant domains at each leaflet contributed by their respective compositions is central to understanding cellular signalling and membrane function.

Using a combination of membrane impermeable lipases, lipidomics and chemical modifications several studies have demonstrated that lipid species such as Sphingomyelin (SM) and Phosphatidylcholine (PC) are predominantly localized to the outer leaflet of the PM whereas Phosphatidylserine (PS) and Phosphatidylethanolamine (PE) are enriched at the inner leaflet^12–15^. Lipids also display a varying degree of unsaturation in their acyl chains associated with leaflet location. While asymmetry of lipid distribution across the leaflets of the bilayer is a fundamental feature prevalent across eukarya and to a major extent in bacteria, the role of this asymmetry is poorly understood^16,17^. The increasing degree of asymmetry from the endoplasmic reticulum (ER) to PM and the consequent membrane lipid environment is suggested to aid in sorting/harbouring proteins of specific kinds at each organelle^3,14^. Functional activity of proteins is also predicted to be affected by the lateral pressure profiles created in synthetic asymmetric membranes^18^. In animals the loss of asymmetry and resultant exposure of PS at the outer leaflet, is a cue to recruit macrophages to recognize and remove these apoptotic cells or recruit factors involved in blood clotting^19,20^. However, investigation of the biophysical consequences of an actively maintained asymmetry at the plasma membrane in living cells and their signalling has been limited. Simulations and limited experiments mainly in red blood cells thus far, indicate that the outer leaflet is more ordered than the inner leaflet but a localized inner leaflet based reporter and measurements are missing from the field^14,21^.

Lipid environments can be explored using a variety of reporters that measure local viscosity, polarity or levels of a particular species^22,23^. One category of such reporters are solvatochromic molecules which respond to the local lipid packing (density) or solvent penetration. This results in a red-shifted emission in liquid disordered (*l_d_*) phases due to its higher local polarity and hydration compared to liquid ordered (*l_o_*) phases in a membrane^22,24^. Probes such as Laurdan (Fig 1a) and its derivatives are well known examples^25,26^. However, since these probes redistribute to all membranes of the cells, the task of characterizing a single membrane is technically challenging, let alone a single leaflet.

**Figure 1:**
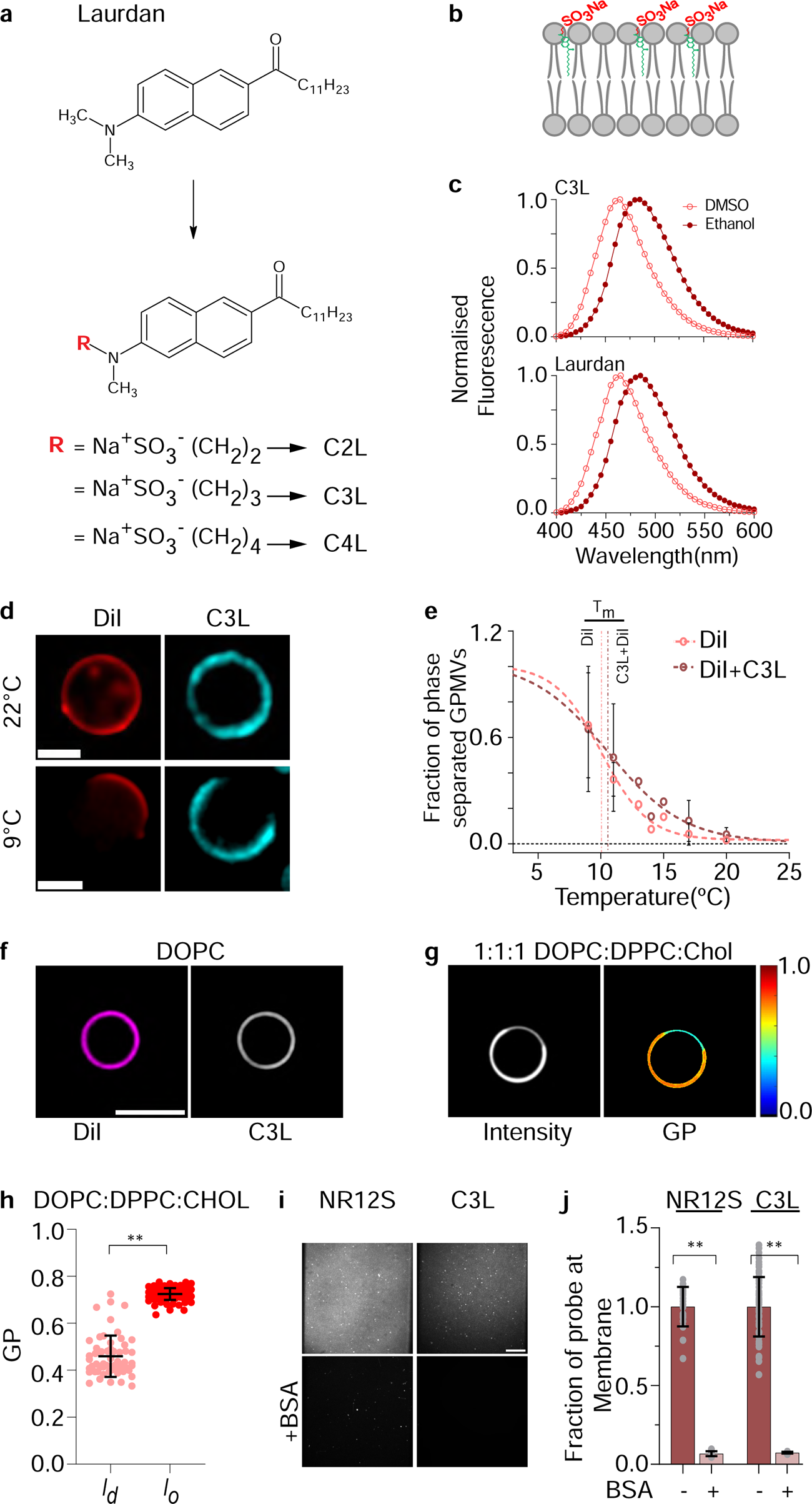
C3L is a *lo* domain preferring solvatochromic reporter. (a) Structures of C(X)L molecules derived from the Laurdan template. (b) Schematic indicating expected distribution and orientation of C3L in the plasma membrane bilayer. (c) Emission spectra of Laurdan(top) and C3L(bottom) in DMSO and Ethanol at λ(ex)= 350nm. (d) Confocal images of GPMVs labelled with C3L(blue) and liquid disordered domain marker Fast DiI(red) at 9°C where they are phase separated and at 22°C where they are homogenous. Scale=5µm. (e) Graph showing the fraction of phase separated GPMVs labelled with C3L (Dotted brown) or C3L with Fast DiI (Dotted pink) at the indicated temperatures. Tm is the temperature at which 50% of the GPMVs are phase separated. Each data point is obtained from at least 90 vesicles. (f) Confocal images of GUVs showing fluorescence emission of Fast DiI (left) and C3L(right) from the same vesicle at 25°C, prepared from DOPC. (g) Confocal images of total intensity (left) and Generalised Polarisation (GP) (right) from C3L labelled DOPC or 1:1:1 DOPC:DPPC:Chol GUVs maintained at 25°C. (h) Graph showing GP values derived from *lo* and *ld* regions C3L-labelled 1:1:1 DOPC:DPPC:Chol vesicles at 25°C. Each dataset is obtained from at least 25 vesicles. (i) TIRF images of 98:2 DOPC:[DGS-NTA(Ni^2+^)] bilayers labelled with NR12S or C3L prior to and after BSA extraction. (j) Graph showing the quantification of amount of probe in the bilayers before (-) or after (+) back extraction with BSA. Data is plotted as mean±s.d. p values determined from unpaired t test or Mann-Whitney test, where ** indicates p<0.005. Unless mentioned, Scale=10µm.

A few solvatochromic reporters such as Di-4-ANEPPDHQ (Di4) and NR12S are localised to the plasma membrane, specifically to its outer leaflet^14,27^. While these have proven to be extremely important in exploring plasma membrane biology they limit the investigation to the outer leaflet. By microinjection into cells, Di4 has been employed to investigate the inner leaflet organization which still limits the capacity to look at both leaflets in the same cell using the same probe because once injected Di4 redistributes to all intracellular membranes^14^.

Here, we report the design and characterisation of a set of probes based on Laurdan to explore the lipid environment of the PM, and specifically the inner leaflet. These probes deploy the hydrophobic aliphatic moiety of Laurdan to anchor the solvatochromic fluorophore into the membrane leaflet and a zwitterionic sulphonate group like that present in NR12S to constrain it to the membrane bilayer (Fig. 1a, b). We find that the probe serves as a substrate for flippases and scramblases at the cell surface. After addition from the outside, this facilitates its redistribution to the inner leaflet, rendering a substantial fraction resistant to back extraction to reveal organization at the inner leaflet.

With the specific probe C3L characterized here, we find that the environment at the inner leaflet is significantly disordered compared to the outer leaflets of cells. We also utilize the properties of the probe to examine the outer and inner leaflet in three conditions in the living cell: (i) where the lipid composition of the cells are specifically perturbed, (ii) modulated during mesenchymal to epithelial transition^28^, or (iii) where local protein scaffolds are constructed such as at focal adhesions or artificially crosslinked protein patches. Our results suggest that the lipid ordering at each leaflet is established mainly by the composition of lipid species, and is locally modulated by the protein scaffolds.

## RESULTS

### Design of a solvatochromic membrane probe for detecting ordered domains

Utilising Laurdan as a template, we designed a series of solvatochromic molecules with a variable number (2-4) of CH2 units as a spacer to link the membrane permeant solvatochromic group of Laurdan to a zwitterionic SO3-Na+ as in NR12S (Fig. 1a; Suppl Text). The zwitterionic group is expected to restrict the probe to the hydrophilic head group layer of the phospholipids and the long hydrophobic alkyl chain to position and stabilise it in the bilayer while the aromatic fluorochrome of Laurdan (see Fig. 1b) allows the molecule to ‘sense’ solvent penetration into hydrophobic region of the bilayer. The synthesis of these probes will be described separately, elsewhere.

### Properties of membrane incorporated C3L

To study their solvatochromic properties, we measured the emission spectra of C(X)L in solvents of differing polarities. We find that all the Laurdan derivatives have overlapping emission spectra that became more red-shifted with increasing solvent polarity (**Fig. 1c, Suppl Fig. S1a**). This indicated that C(X)L structure is solvatochromic and may be used to visualise the extent of solvent penetration in a lipid bilayer. We proceeded with C3L for the remaining study with solvatochromic properties as sensitive as any of the other analogues.

To capture the behaviour of C3L in a complex lipid mix with different degrees of solvent penetration, we prepared phase separating Giant Plasma Membrane Vesicles (GPMVs) from cells^29^ labelled with C3L and the *l_d_* domain marker, Fast DiI. To induce phase-separation, the GPMVs were subjected to a stepwise reduction in temperature from 22°C to 9°C. At 22°C barely any of the GPMVs exhibited detectable phase-separation and the fluorescence distributions of the two probes were coincident (**Fig. 1d**). Upon lowering the temperature to 9°C, around 80% of the GPMVs exhibited clear phase-separation (**Fig. 1d, e**). The phase separated regions enriched in Fast DiI, showed a marked anti-correlation with C3L fluorescence as noted by the partition coefficient of the probes in the respective domains (**Table S1, Suppl Text**), indicating that in a phase separated lipid bilayer C3L has a preference for *l_o_* domains compared to *l_d_* domains. To rule out if incorporation of the probe alters biophysical properties of the membrane bilayer, we determined the transition temperature (T_m_) of a population of GPMVs labelled with C3L and Fast DiI or Fast DiI alone. We find that the percentage of separated GPMVs is similar between the two at all temperatures, *i.e.* the T_m_ is is unperturbed by C3L (**Fig. 1e**). These results indicate that C3L is a solvatochromic probe with a preference for *l_o_* domains and at the concentrations used does not significantly alter the phase segregation characteristics of GPMVs.

To study whether the solvatochromic properties of C3L can be used to distinguish between liquid-ordered (*l_o_*) and disordered (*l_d_*) phases in a synthetic membrane, we prepared Giant Unilamellar Vesicles (GUVs) from two kinds of lipid mixtures, one from pure 1,2-Dioleoyl-sn-glycero-3-phosphocholine(DOPC) and another from an equimolar mixture of DOPC, dipalmitoylphosphatidylcholine (DPPC) and Cholesterol (Chol)^30^. The former are expected to exhibit a homogenous *l_d_* phase at 24°C whereas the latter phase separate into *l_o_* and *l_d_* phases^24^. Fast DiI staining showed that GUVs made from DOPC exhibit a uniformly labelled distribution of Fast DiI and consequently a homogenous *l_d_* phase whereas the DOPC:DPPC:Chol(1:1:1) mixture exhibits two distinct phases at 25°C marked by segregation of Fast DiI (**Fig. 1f**, **Suppl Fig. S1b**). Consistent with observations on GPMVs, C3L distributed to domains poorly labelled by the Fast DiI in the GUVs (**Suppl. Fig. S1b**) indicating that C3L partitions into the *l_o_* phase, similar to its precursor, Laurdan (**Suppl Fig. S1c**).

We next monitored the solvatochromic shift of C3L in these membranes, characterized by its Generalized Polarization (GP), a metric for the extent of solvent relaxation that results in its emission maxima shifting to longer wavelengths. GP is the normalised ratiometric measurement of fluorescence emission of C3L in two emission bandwidths, one centred at 440 nm corresponding to a more ordered phase and the other at 490 nm, the more red-shifted wavelength corresponding to disordered phase (see Methods)^31^. These are identical to the spectral bandwidths used for previous measurements of GP of Laurdan and chosen owing to the overlapping emission spectra. We measured the GP reported by C3L in the *l_o_* and *l_d_* domain of the phase separating vesicles labelled with only C3L (**Fig. 1g**). We find the *l_o_* domains in the GUVs have a GP of 0.72±0.02 while the *l_d_* domains exhibit a GP of 0.46±0.08 (**Fig. 1h**). In Laurdan-labelled GUVs, the *l_o_* and *l_d_* phases also differ by 0.28±0.1, similar to C3L (**Suppl Fig. S1d**). Together with the data derived from GPMVs, these data show that C3L serves as a suitable solvatochromic probe that partitions into *l_o_* domains in synthetic and PM-derived membranes and therefore can serve as a reporter to assess lipid packing with a significantly high dynamic range.

To address the leaflet distribution of C3L once it is added to synthetic membranes, we labelled supported lipid bilayers composed of 98:2 DOPC:DGS-NTA(Ni^2+^) with C3L or NR12S that is incorporated into the outer leaflet^27,32^ (**Fig. 1i**). We employed BSA based back extraction, a method used to remove fluorescent lipids which remain confined to the outer leaflet of the cell membrane^33^ . This resulted in the quantitative extraction of both probes from the bilayers (**Fig.1i, j**). Nevertheless, labelled His-tagged protein that binds to the Ni^2+^-NTA lipids and added to the bilayers prior to back extraction, was quantitatively retained, consistent with the supported bilayers remaining intact after back extraction with BSA (**Suppl Fig. S1e, f**). These observations confirm that C3L and NR12S are efficiently extracted by BSA extraction after their incorporation into membranes of synthetic bilayers, consistent with an outer leaflet disposition.

### C3L distributes across both leaflets in the plasma membrane of cells

We next labelled cells with Laurdan, C3L or NR12S and assessed their sub-cellular distribution in living cells in a confocal microscope (**Fig. 2a**). While Laurdan extensively labelled the cell surface as well intra-cellular compartments (**Fig. 2a**) as previously reported^34^, C3L and NR12S fluorescence were restricted to the plasma membrane under similar labelling conditions. Cells labelled with solvatochromic probes, NR12S or C3L, were imaged before and after exposing the incorporated probes to BSA based back extraction (**Fig. 2b**). While NR12S was completely extracted, C3L was only partially extracted from the cell membrane (**Suppl Fig. S2a**). We reasoned that given the extraction should be limited to the outer leaflet, C3L is either trapped at the plasma membrane by a membrane binder and/or is also distributed to the inner leaflet thereby preventing its complete extraction by the BSA added to the buffer bathing the cells. To address the former possibility Fluorescence Correlation Spectroscopy (FCS) measurements were carried out to determine the diffusion coefficient of C3L in the membrane, both before and after back-extraction. The data shows that although the number of diffusing species is almost halved, the diffusion coefficient and brightness of the mobile species remained unchanged, prior to or post-BSA extraction (**Suppl Fig. S2b, c**), consistent with a highly mobile lipid species^35^. These results do not support the possibility of membrane retention due to association with a slower diffusing membrane molecule such as a membrane protein.

**Figure 2:**
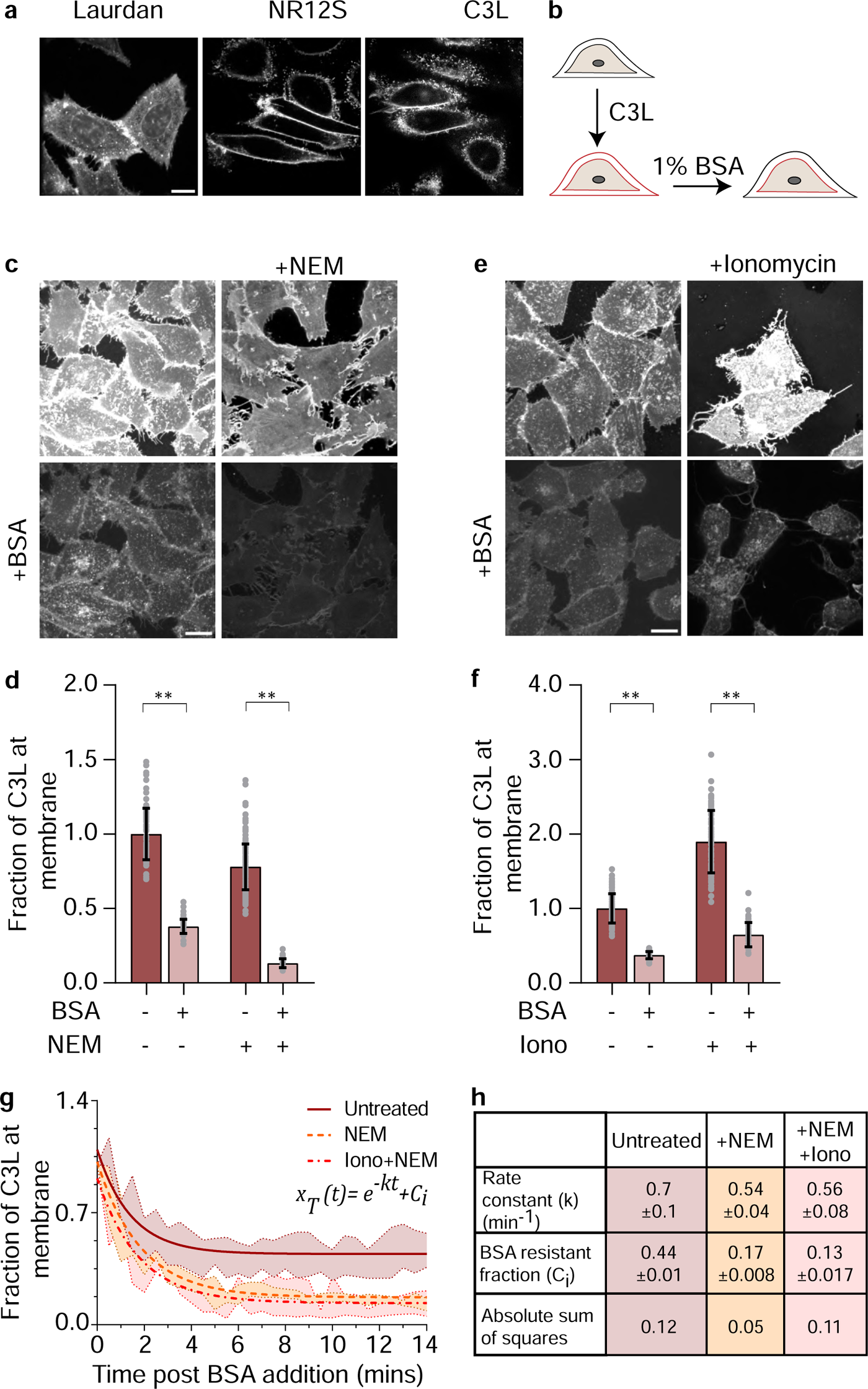
C3L is distributed across the two leaflets of the plasma membrane by flippase and scramblase activity. (a) Confocal images of equatorial plane of cells labelled with 1uM Laurdan, NR12S or C3L showing the sub-cellular distribution of the respective probes. (b) Schematic showing the BSA back extraction assay used for extracting the outer leaflet population of probe in adherent cells. (c-f) Confocal images (c, e) collected from the bottom membrane plane of untreated cells (left panels) or cells treated with 2mM NEM to inhibit flippase activity (c) or 10uM Ionomycin to activate scramblase activity (e), before (top) and after (bottom) back extraction by BSA. Graphs (d, f) show the extent of incorporation (-) and remainder (+) of C3L after extraction with BSA. Each dataset is normalized to the mean intensity of untreated cells labelled with C3L. Data were obtained from from at least 40 cells. (g) Graph shows the time course of fluorescence intensity of C3L remaining at cell surface as a function of exposure to BSA added at t=0 mins (obtained using a CSU22 spinning disk confocal) in untreated cells (-), NEM treated cells (--) and NEM+Ionomycin treated cells (-.-). Each dataset is normalized to its mean starting intensity at t=0mins. Lines indicate fits to single exponential decay where k is the rate constant for the decay in fraction of C3L at the membrane after adding BSA, C_i_ is the BSA resistant fraction after back-extraction. Data were obtained from atleast 20 cells in each condition. (h) Table shows rate constant (k) and BSA resistant fraction (C_i_) obtained from the fit to a single exponential decay shown in panel g. All data is plotted as mean±s.d. p values determined from unpaired t test where ** indicates significance indicated by p<0.005. Scale=10µm.

We next performed a series of experiments to determine the characteristics of the re-distribution of C3L across the two leaflets by varying the concentration of C3L and time after labelling. The data show that C3L distribution rapidly reaches a steady state that is independent of the concentration of the added probe; ∼70% of C3L is back extracted over the range of concentration and residence times examined (**Suppl Fig.S2 d, e**). The consistent back extraction-resistant fraction, together with the ability to quantitatively extract the probe from an artificial bilayer suggested that the redistribution of C3L is not spontaneous. These results strongly suggest that the extraction resistant fraction is maintained by an active cellular process that translocates the probe to the inner leaflet.

### Leaflet distribution of C3L is regulated by flippases and scramblases

The transbilayer distribution of lipids of the PM is actively maintained by a combination of sites of synthesis and the activity of transmembrane proteins such as flippases, floppases and scramblases that redistribute lipids to generate transbilayer asymmetry^2^. To determine if some of the same processes could regulate the distribution of C3L, we perturbed the activity of these proteins. First, the activity of flippases was inhibited by treating cells with 2mM NEM which resulted in PS exposure at the outer leaflet^36^, as confirmed by annexin binding to NEM treated cells (**Suppl Fig. S2f**). In NEM treated cells, not only was the amount of probe incorporated at the membrane reduced but there was a significantly smaller BSA-resistant fraction of the probe (**Fig. 2c, d, Suppl Fig. S2g**). This confirms the role of a NEM-sensitive flipping mechanism in the transfer of the probe to the inner-leaflet.

In a different approach scramblases were activated by treating cells with the Ca^2+^ ionophore, ionomycin^37^. Rapid scrambling of inner leaflet PS amongst other lipid species was highlighted by annexin binding at the outer leaflet (**Suppl Fig. S2f**). Upon ionomycin treatment, cells show a higher labelling efficiency of C3L (**Fig. 2e, f, Suppl Fig. S2h**). This indicates that once activated, scramblases are involved in shuttling C3L across the two leaflets and assisting in the incorporation of the probe.

To understand the dynamics of the re-distribution of the probe between the leaflets once it is incorporated into the membrane, we monitored the fluorescence intensity of the probe at the membrane in real time (**Fig. 2g**). The time dependent evolution of C3L levels in the membrane after BSA addition to extract outer-leaflet localized molecules were fitted to an exponential decay (**Fig. 2g**). The extraction kinetics of the probe followed a single exponential decay with a half time of 1 min, leaving behind a much slower exchanging population at the end of 15 min (**Fig. 2h**). This pointed to a significant fraction of the probe being subjected to rapid back extraction by BSA followed by a very slow exchange phase (see also **Suppl. Fig. 3c**). The kinetics of C3L extraction in NEM or NEM and Ionomycin treated cells exhibited only a very small difference in the rate of back extraction however there was a significant reduction in the BSA-resistant fraction at the end of the extraction (**Fig. 2g, h**). This is consistent with NEM-sensitive flippases playing a major role in transporting C3L to the inner leaflet and basally active scramblases shuttling the probe between the two leaflets. Together these results demonstrate that when added to cells, C3L has a significant inner leaflet population that is driven and maintained by flippase/scramblase activity.

### C3L reports on outer and inner-leaflet membrane order

To measure the heterogeneity in lipid organisation at each leaflet of the plasma membrane of intact cells we first ascertained that C3L incorporation does not systematically influence membrane organisation. We measured the GP of the membrane at different labelling concentrations of C3L (300nM to 2uM; **Supp. Fig S3a**). Empirically, labelling with 1uM C3L in these cells appeared optimal to balance detection sensitivity with minimal perturbation of membrane order. To assess the lipid order of the membrane and especially the inner leaflet, cells were labelled with 1uM C3L and imaged before and after adding BSA in the extracellular buffer at RT on the microscope stage to determine GP. Prior to adding BSA, the GP is relatively high (0.62±0.02). After BSA addition, coincident with the plateauing of the fraction of C3L remaining in the membrane, the GP values also plateau at a significantly lower value (0.4±0.02), albeit at a slightly longer timescale (**Fig 3a, b**), showing no further change until 40 mins (**Suppl Fig. S3c, d**). The GP_T_ (the weighted average of the contribution of the GP of the two leaflets) of the membrane recorded before BSA extraction is significantly higher than GP_i_ (GP of the inner leaflet) recorded when no further changes in intensity are seen after BSA-extraction. To rule out that the differences between GP_T_ and GP_i_ is not caused by BSA extraction, GP_T_ was assessed before and after BSA-extraction. Although there is a negligible increase of 0.02±0.08 in GP_T_ between the BSA pre-exposed vs unexposed cells, the difference in GP_T_ and GP_i_ of 0.2±0.04 is an order of magnitude larger (**Suppl Fig. S3b**). This is likely driven by the asymmetric composition of the two leaflets. From these datasets we concluded that we have a method to measure the GP of the inner leaflet (GP_i_) using C3L as a reporter, near simultaneously with that of the bilayer (GP_T_).

**Figure 3:**
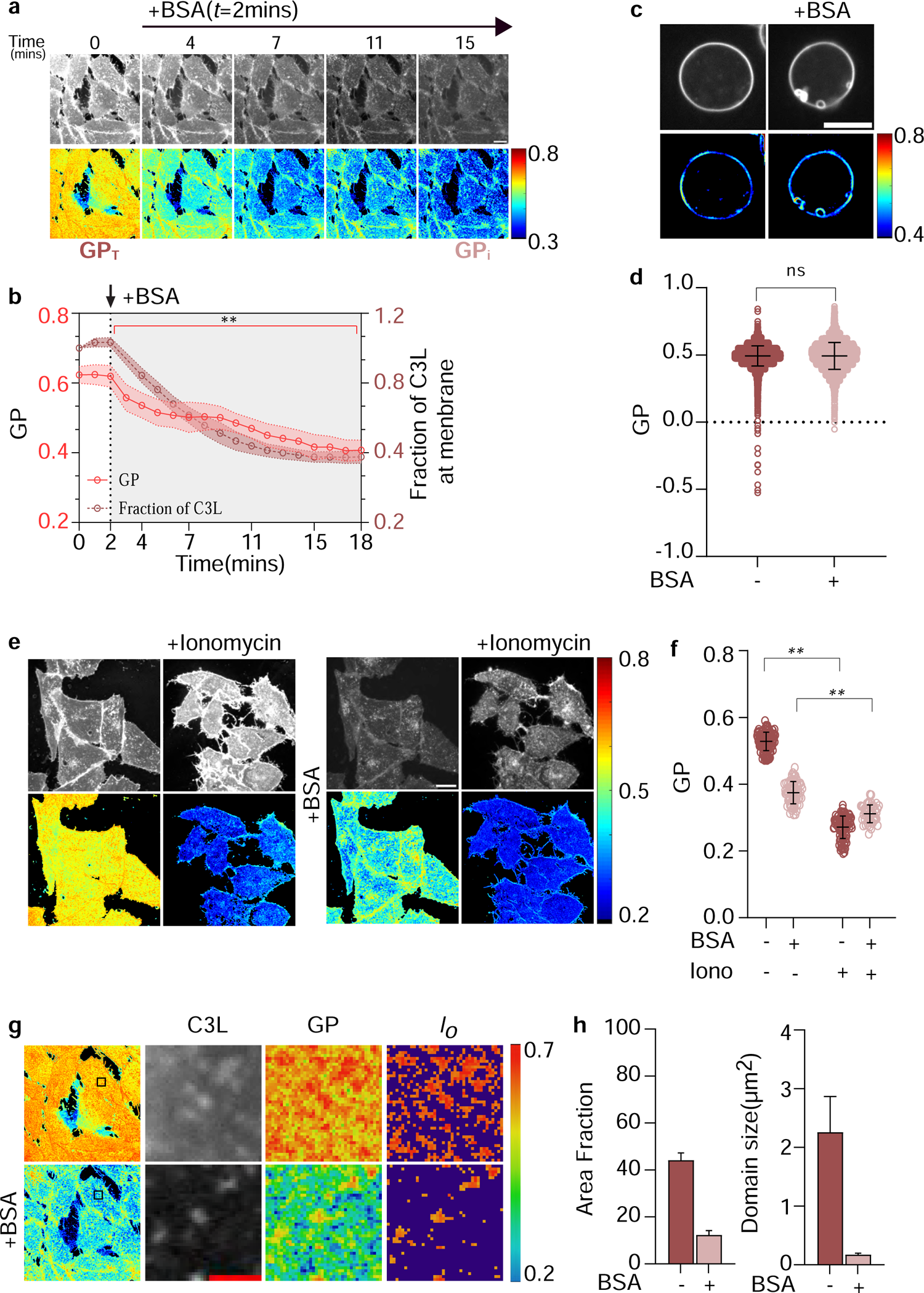
The nature of the inner leaflet revealed after BSA extraction of C3L incorporated at the cell surface. (a) Confocal montages of fluorescence emission intensity (top) and GP (bottom) distribution collected from the bottom plane of C3L-labeled cells subjected to BSA extraction, beginning at t=2mins. (b) Graph depicts simultaneous intensity (right Y axis) and GP values (left Y axis) determined from the time course of BSA extraction of C3L labelled cells in panel a. At t=0mins, GP_T_ is the weighted average of the GP of the bilayer, and at t=15mins, GP_i_ is the GP of the inner leaflet. Data quantified from 35 cells (c) Confocal images of fluorescence emission intensity of C3L labelled GPMVs (top) and their corresponding GP maps (bottom) before (left) and after(right) exposure to BSA extraction for 15mins (d) Graph shows GP values determined from the GPMVs before and after BSA extraction as shown in panel c. Data taken from atleast 40 vesicles. (e) Confocal images of fluorescence intensity and GP from C3L-labelled cells before or after BSA extraction as indicated. Cells were either untreated (left) or treated with ionomycin (10µM) at 37°C (right). (f) Graph shows GP values obtained from the cells in panel e. Data are taken from 20 by 20ROIs from atleast 40 cells. (g) 4.4µM by 4.4µM patches taken from cell surface in panel (a). The images indicate intensity of C3L (left), corresponding GP map (middle), and the thresholded GP map showing regions representing domains with GP>0.65 (GPmean(GPMV)+2SD) for top, GP>0.57(GPmean(GPMV)+1SD for bottom) before (top) and after (bottom) BSA-back extraction. Scale= 2µm. (h) Graphs showing mean±s.e.m of area fraction of the membrane covered higher order domains and their sizes from the thresholded images of GP_T_ and GP_i_ shown in panel g. Data are taken from 4.4µm by 4.4µm ROIs from 22 cells. Unless mentioned data is plotted as mean±s.d . *p* values determined from unpaired t test where ns indicates no significant difference and ** indicates significance indicated by p<0.005. Scale=10µm.

To ascertain whether the difference in GP_T_ and GP_i_ is indeed due to compositional differences between the two leaflets, we reduced this compositional asymmetry by two methods. First, GPMVs were made from cells labelled with C3L and GP was measured before and after back extraction (**Fig. 3c**). GPMVs are known to have symmetric lipid composition^38^, and consistent with our hypothesis GP measurements with C3L showed that there was no difference between GP_T_ and GP_i_, (0.49±0.07) in GPMVs; **Fig. 3d**). Second, GP_T_ and GP_i_ were measured in adherent cells treated with ionomycin where the activation of scramblases should rapidly equalize the lipid composition across the bilayer (**Fig. 3e**). The difference between GP_T_ and GP_i_ was lost in these ionomycin treated cells as well (**Fig. 3f**). Taken together these data show that the differences between GP_T_ and GP_i_ in unperturbed cells upon back extraction, are a result of intrinsic compositional differences between the two leaflets.

Using the distribution of GP values from GPMVs in **Fig 3d**, we quantified the heterogeneities at the membrane and the inner leaflet assuming GP>(GP_mean_)_GPMV_ to indicate ordered domains and GP<(GP_mean_)_GPMV_ disordered. Given GP_T_ is significantly higher, we thresholded GP values higher than mean+2SD from the GPMV distribution to specify more ordered regions at the membrane (**Fig. 3g**). We find that these domains are present in ∼45% of the membrane and have size of ∼2.3µm^2^ (**Fig. 3h**). Domains with equivalent order were not found at the inner leaflet. Instead in the GP_i_ distribution, we do find domains with higher order than the surrounding, with a GP value higher than mean+1SD of the GPMV distribution (**Fig. 3h**). With these parameters we find higher order regions in ∼10% of the leaflet that have domains sizes ∼0.3μm^2^ and often originate from regions associated with structures that concentrate C3L (**Fig. 3h**).

### Lipid class-specific effects on leaflet organisation at a cell surface

As suggested above, the lipid composition of each leaflet appears to play a predominant role in determining the GP of the leaflet. Therefore, we reasoned that by altering the levels of specific lipid classes located at each leaflet we would be able to assess their influence on lipid packing. This could give us a broader understanding about the mechanism of setting up transbilayer asymmetry in lipid ordering.

We first examined the role of sphingomyelin (SM), a predominantly outer leaflet localised lipid in influencing the GP of the membrane. A fungal toxin Fumonisin B1 that interferes with the synthesis of ceramides was used to deplete sphingomyelin levels in the cell and therefore reduces SM levels at the membrane^39^. In agreement with its predominantly outer leaflet localisation, SM depletion only reduced GP_T_ but had no effect on GP_i_ (**Fig.4a, Suppl Fig. S4a**). Given that GP_T_ is a weighted average of the two leaflets, we estimated the GP of the outer leaflet (GP_o_) from values of GP_T_, GP_i_ and the respective fractions of probe in each leaflet (see methods for further explanation). Plotting the frequency histogram of GP_T_, GP_i_ and GP_o_ showed a reduction (leftward shift) of the mean GP_T_ and GP_o_ to lower GP values with no effect in the mean of GP_i_ (**Fig. 4b**) indicating SM regulates order only at the outer leaflet.

**Figure 4:**
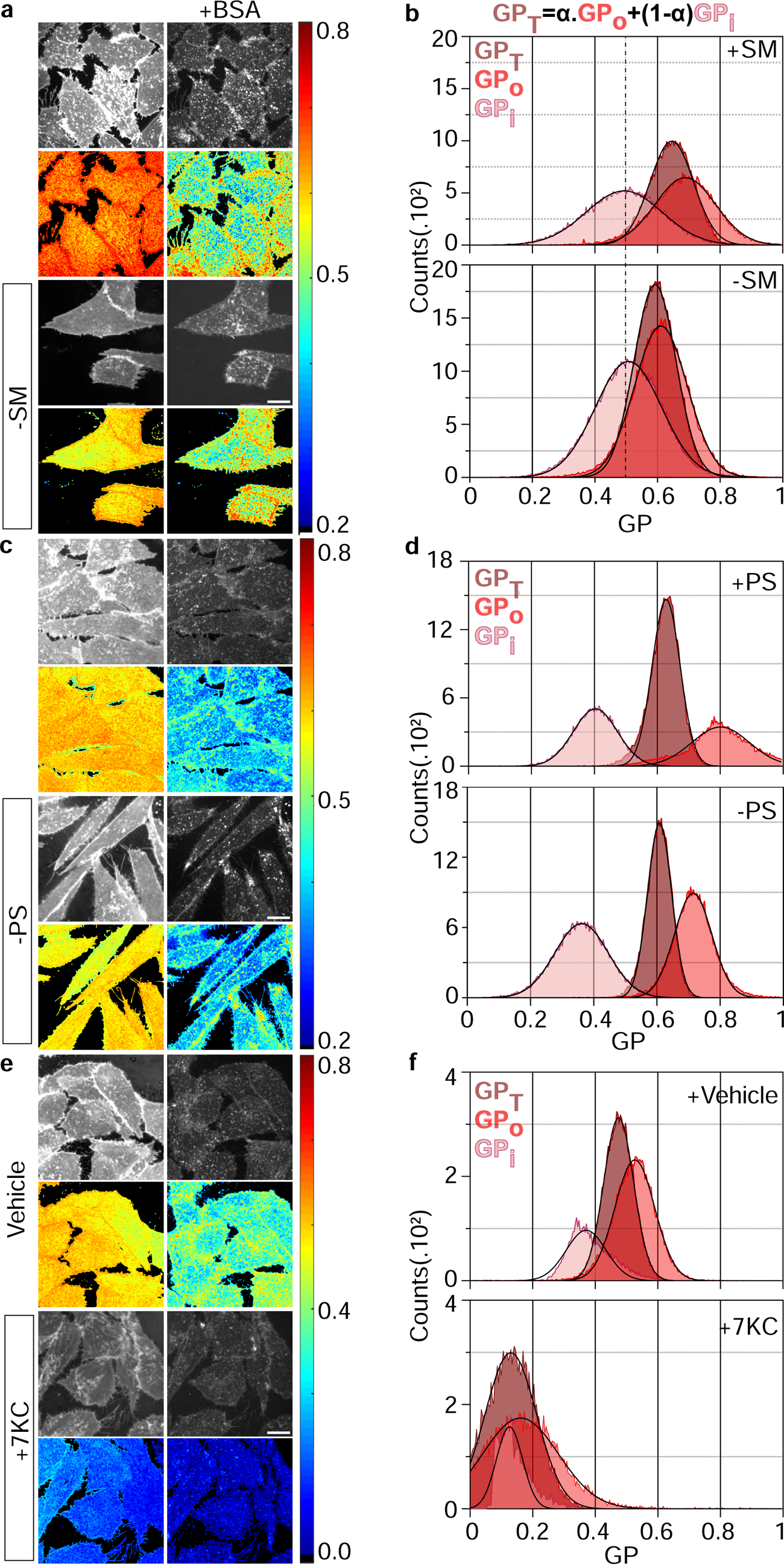
Specific lipid perturbations selectively modulate the GP of each leaflet of the membrane. (a-d) Confocal images (a, c) of fluorescence intensity and GP of cells depleted of the indicated lipids (-SM, sphingomyelin; -PS, phosphatidylserine) show changes induced by indicated lipid perturbations. (b, d) Corresponding frequency distributions of the GP values for GP_T_, GP_i_ and GP_o_ from the same cells determined as described in Methods, and shown in the right. Data obtained from ≥15 cells in each condition. (e, f) Confocal images of fluorescence intensity and GP (e) of cells either treated with vehicle (0.38mM MBCD + Ethanol) or vehicle + 7-ketocholesterol (7KC) and their corresponding frequency distributions (f) of GP values for GP_T_, GP_i_ and GP_o_. Data were obtained from ≥ 8 cells per condition, Scale=10µm.

We next examined the role of phosphatidylserine (PS), an inner leaflet component, using cells which are defective in the PSS II enzyme and have a compromised PS metabolism^40^. In these cells PS levels are depleted by ethanolamine deprivation. Although PS is expected to be predominantly inner leaflet localised, its depletion reduced both GP_T_ and GP_i_ (**Fig. 4c, Suppl Fig. S4b**). The frequency histogram of GP_T_, GP_i_ and GP_o_ showed a corresponding left shift of mean GP_T_, GP_O_ and GP_i._ (**Fig. 4d**) indicating PS regulates order both at the inner and outer leaflet though itself is predominantly present at the inner leaflet.

Finally, we examined the role of cholesterol, given its ability to modulate the transition of both gel and fluid phases to liquid ordered phase. While the plasma membrane contains ∼40% cholesterol, its precise leaflet distribution remains debated^41^. To disrupt ordered domains, we introduced the cholesterol derivative, 7-Ketocholesterol (7KC), using low concentration cyclodextrin as a vehicle^42^. Introducing 7KC in the membrane drastically reduced the order of both leaflets as indicated by the dramatic leftward shift in the frequency histogram of GP_T_, GP_i_ and GP_o_ (**Fig.4e, f Suppl Fig. S4c**). This clearly demonstrates that cholesterol plays an important role at both leaflets.

### Probing lipid packing in different cellular contexts

Using the tools and protocols developed here, we explored three different cellular contexts where lipid packing is likely to be specifically altered at the level of the two leaflets. These are i) a mesenchymal to epithelial cell transition, ii) the generation of membrane localised sub-cellular signalling structures such as focal adhesions, and iii) protein clustering.

During epithelial morphogenesis, polarised epithelial cells accumulate saturated and longer sphingolipid species along with an increase in glycosphingolipid and cholesterol content^28^. These cells are characterized by the generation of distinct domains, apical and basolateral, maintained by protein dense structures called tight junctions (TJs)^43^. We reasoned that these changes should be reflected in alterations in lipid packing in the bilayer, as well as in individual leaflets. To explore the consequences of these changes, MDCK cells in their mesenchymal (**Fig 5a**) and epithelial (**Fig 5c**) states were labelled with C3L and imaged prior to and after BSA-back extraction (**Fig. 5b, 5d**). Three salient observations stand out. First, there is significant increase in the GP_T_ in the epithelial state compared to the mesenchymal state. Second, GP_i_ in both mesenchymal and epithelial states is indistinguishable, being significantly lower than their respective GP_T_ (**Fig.5b, d, e**). The increase in the GP_T_ in the epithelial state must therefore come from a tighter lipid-packing in the outer leaflet. Thirdly, in the epithelial cells as the membrane is traversed along the basal (at x=0) to the apical (at x=1) axis, there is a significant increase in GP_T_ at the apical domain of the cell (again without any significant increase in GP_i_) suggesting that these changes must be due to higher GP_o_ at the apical domain (**Fig 5e, f, g**). These observations suggest that during mesenchymal to epithelial morphogenesis, a much tighter lipid packing is developed mainly in the outer leaflet of epithelial cells.

**Figure 5:**
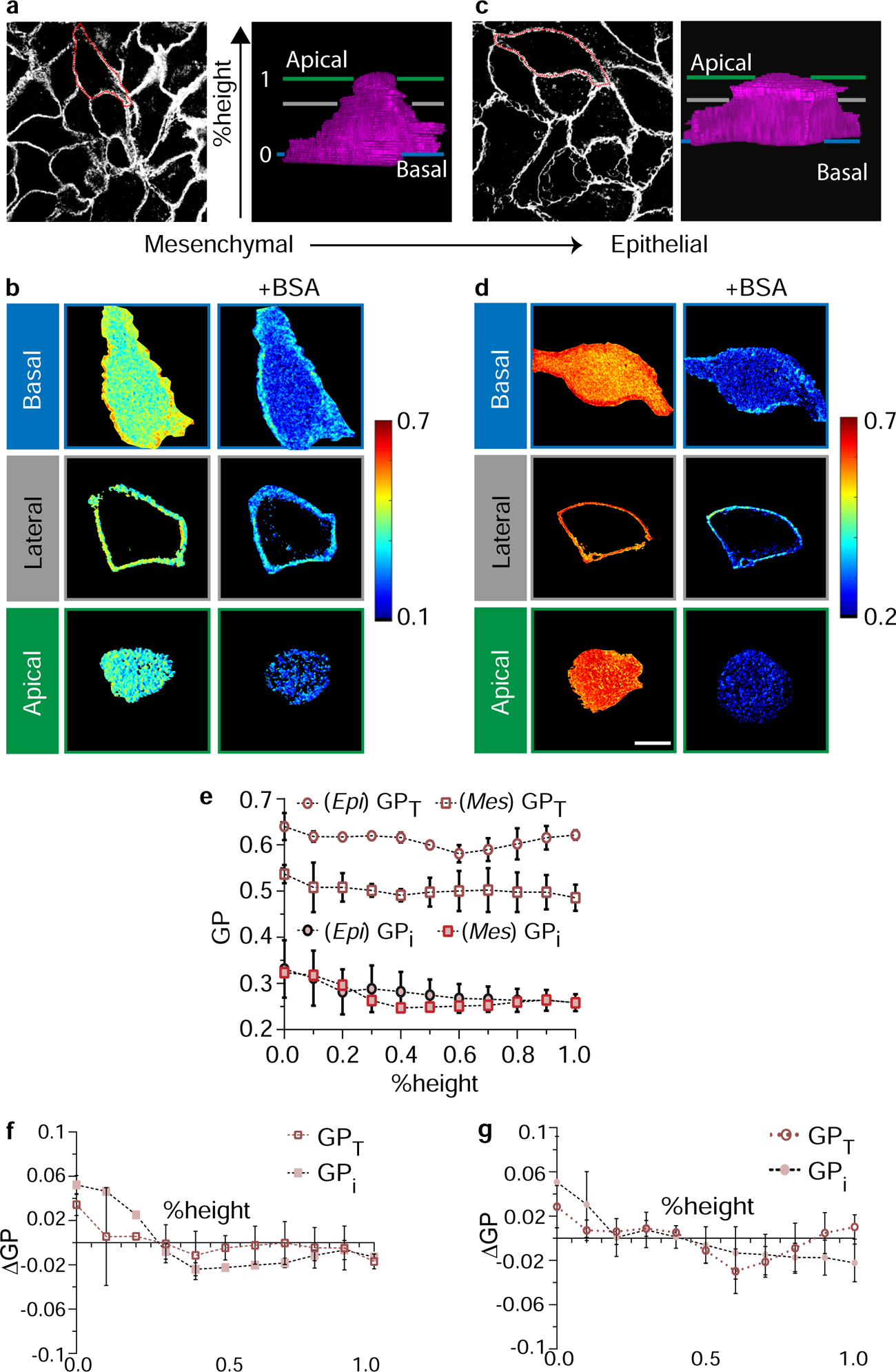
Epithelial cells show changes in membrane lipid order with differentiation. (a) Confocal cross-section of the cells (left) and 3D reconstruction (right, magenta, from membrane signal acquired using C3L labelling of the marked MDCK cell 1 day after plating, in mesenchymal state, with lines representing basal (blue), lateral (gray) and apical (green) domains of the cell. (b) Confocal GP images of the cell in right of panel (a) from the cross section at basal, lateral and apical domain before(top) and after (bottom) BSA back extraction. (c) Confocal cross-section of the epithelial sheet (left) and 3D reconstruction (right, magenta from membrane signal acquired using C3L labelling) of the marked MDCK cell, 5 days post 100% confluency, in epithelial state, with lines representing basal (blue), lateral (gray) and apical (green) domains of the cell. (d) Confocal GP images of the cell in the right of panel (c) from the cross section at basal, lateral and apical domain before(top) and after (bottom) BSA back extraction. (e) Graph showing absolute values of GP_T_ and GP_i_ of the cells in epithelial (epi) and mesenchymal (mes) states along the height of the cell (basal=0, apical=1). (f) Graph showing the difference in GP(GP-GP_mean_) along the height of the cell (basal=0, apical=1) w.r.t to the mean GP of the cells in mesenchymal state from 25 cells from panel b. (g) Graph showing the difference in GP (GP-GP_mean_) along the height of the cell (basal=0, apical=1) w.r.t to the mean GP of the cells in epithelial state from 90 cells from panel d. Scale=10µm.

The results obtained thus far indicate that the outer leaflet is a much more tightly packed environment compared to the inner leaflet. We next wondered how proteins and their complexes are accommodated and affect or get affected by the lipid environment. Focal adhesions (FA) form large visible structures at the interface of the plasma membrane and the extracellular matrix. These long lived complexes are formed as a cell senses and exerts force on the ECM during migration. FAs consist of a number of transmembrane and peripheral proteins such integrins, vinculin and paxillin that eventually recruit the cortical cytoskeleton and are enriched in GPI-anchored protein nanoclusters^44^. We marked the FA complexes using mCherry tagged Paxillin and measured GP_T_, GP_i_ and GP_o_ around these complexes in cells co labelled with C3L (**Fig 6a-c**). Plotting the frequency histograms reveals that GP_o_ is unchanged at the focal adhesions in comparison to the rest of the cell surface (**Fig 6d**). However, the change in FA-membrane is mainly due to the stark increase of GP_i_ of the inner leaflet at the FA when compared to the regions without focal adhesions as indicated by the right shift of the mean GP_i_ at the FAs (**Fig. 6d**). These results suggest that signalling active hubs can reorder the inner leaflet.

**Figure 6:**
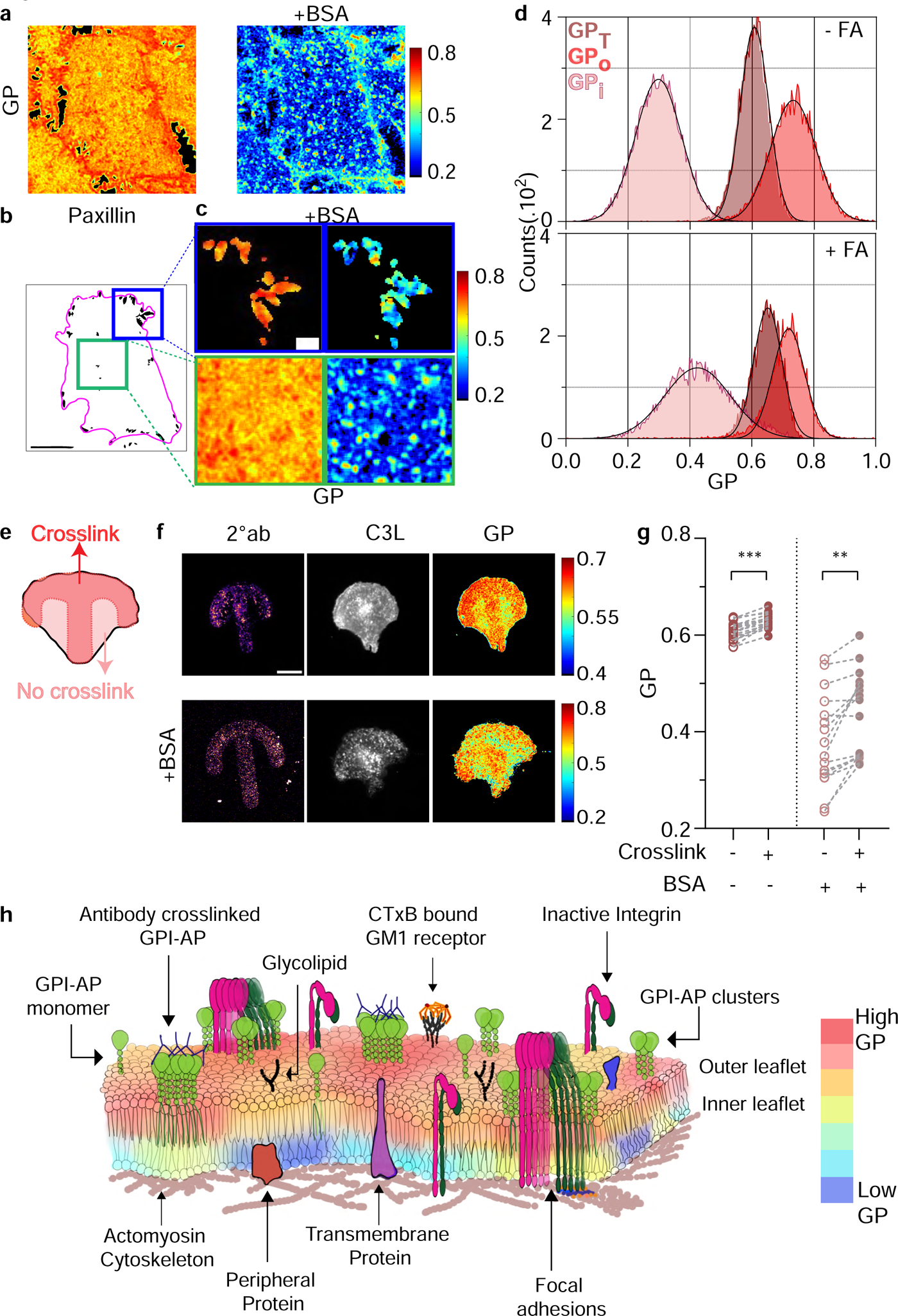
Protein crosslinking generates increases in lipid ordering predominantly at the inner leaflet. (a-d) C3L-labelled cells expressing mCherry-Paxillin were imaged on a confocal microscope and the GP map of the membrane was determined, before (a) and after BSA extraction. (b) mCherry-Paxillin labelled focal adhesions in the same cell were simultaneously imaged (c) and the GP map of regions strongly labelled by paxillin (top; Green ROI in c) or devoid of paxillin (bottom; red ROI in c) are shown corresponding to the boxed section, before (left) and after (right) BSA extraction. Scale=2µm. d) Graph shows the frequency distribution of GP_T_ (brown), GP_i_ (pale pink), and the calculated GP_o_ (red) from focal adhesions (bottom) compared to regions devoid of adhesions (top) quantified from from 4.4µm by 4.4µm across 24 cells (e) Schematic shows a cell plated on a patterned coverslip coated with a crosslinking antibody against a membrane protein (darker shade) designed to compare the GP of the membrane regions overlapping with antibody signal with that of the non-crosslinked antibody-free membrane patches (lighter colour). f) Confocal images of AF647 tagged 2°*ab* covalently attached to patterned coverslips (left), C3L fluorescence emission intensity images (middle) and GP maps (right) of cells expressing folate receptor plated on a pattern coated with secondary followed by primary antibodies against the folate receptor before (top) and after BSA extraction (bottom). g) Graph showing GP values from the regions located on or off the patterns (+/-2°*ab,* before and after BSA extraction. Each datapoint is from a single cell with corresponding points on and outside the pattern of the same cell. Data is taken from ≥16 cells. All data is plotted as mean±s.d, *p* values determined from paired Wilcoxon non-parametric test where ** indicates significance indicated by p<0.005, *** indicates significance indicated by p<0.0001. Unless mentioned, Scale=10µm (h) Graphical summary of the transbilayer leaflet organization. LUT indicates GP values. The outer leaflet is predominantly ordered as indicated by the relatively homogenous GP in red and organge shades. The inner leaflet is predominantly disordered as indicated by the GP in different shades of blue. Large protein scaffolds such as focal adhesions and crosslinked GPI-APs or crosslinked gangliosides such as the GM1 receptor induce negligible increase in ordering of the outer leaflet but a large increase in the ordering of the inner leaflet, indicated by change of GP to yellow locally around these scaffolds at the inner leaflet.

We next asked whether the outer leaflet GPI-anchored protein clusters which form hot spots around FAs could act as a mechanism to generate regions of higher packing at the inner leaflet^45^. We used coverslips with pre-fabricated patterns of activated patches etched on them. We coated these coverslips with secondary antibody(2°*ab)* followed by primary antibody, Mov19 raised against the folate receptor, an outer-leaflet localized GPI-anchored protein and plated cells on them (**Fig. 6e**). As previously shown CHO cells over-expressing the folate receptor spread on anti-folate receptor antibody coated surfaces bypass the need for integrin receptor activation^5^. We find that the membrane order before and after extraction shows an increase along the coated surfaces where the protein would be crosslinked due to antibody binding at the surface. This increase is accompanied predominantly with increases in the inner leaflet (**Fig. 6f, g**). GM1 receptor crosslinked by binding to the pentavalent Cholera Toxin B subunit immobilized on a patterned cover-slip (**Suppl Fig. S5a, b**) also results in similar spatial correlations^46^. Given the high value of GP_o_, a local increase in GP_i_ around these protein scaffolds suggests these multi-valent protein complexes create local ordered domains by altering the inner leaflet. These spatial signatures of ordering at each leaflet and local transbilayer coupling can be read out directly with the reporter C3L developed here.

## DISCUSSION

In this report, we designed and characterised a solvatochromic reporter, by adding a sulphonate group with a 3 carbon linker to the tertiary amine in Laurdan (**Fig.1a**), to image lipid organisation of the two leaflets of the plasma membrane bilayer^3,8^. C3L, like Laurdan, redistributes to *l_o_* domains in lipid mixtures that exhibit *l_o_-l_d_* phase-separation without influencing the phase transition temperature (T_m_) of GPMVs (**Fig 1d-e**). Unlike Laurdan, C3L with its sulphonate group does not passively flow out of the membrane bilayer due to its ability to interact with lipid head groups, and its leaflet distribution is likely regulated by membrane enzymes, flippases/floppases and scramblases^2,47^. At this time, we are not aware of the molecular identity of the flippases/floppases and scramblases. Type IV P-type ATPases (P4-ATPases), such as ATP8B1 which recognise PC in mammalian systems could be involved in flipping C3L to the inner leaflet due to structure of the hydrophilic group in C3L, and its redistribution to the outer leaflet could be maintained by non-specific basal activity of GPCR like scramblases in the plasma membrane^48,49^. Consistent with these possibilities, the inhibition of flippases (by NEM) or activation of scramblases (by increasing intracellular Ca^+2^) exhibit corresponding increases in BSA extractable fractions (**Fig. 2**). Together these properties and the capacity of C3L to be localized to the inner-leaflet, make it a suitable probe to detect and visualize ordered membrane domains with high sensitivity at both leaflets.

Using Generalised Polarisation(GP) of C3L as the quantitative parameter to estimate the degree of lipid packing in different membrane phases, we explored the packing of lipids of the two leaflets taking cues from GUVs of known composition. GP of C3L in the *l_o_* phase of the phase-separated DOPC/DPPC/Chol GUVs is 0.72±0.02 and that of the *l_d_* phase is 0.46±0.08(**Fig. 1f, g**). By comparison, in cell membranes we see that the GP_T_ (weighted GP of the two leaflets) is 0.62±0.02, and the GP_i,_ (GP of the inner leaflet) measured after back extraction with BSA is 0.4±0.02 (**Fig. 3a**), thus, the GP_o_ (GP of the outer leaflet) is even higher, ∼0.65, close to that of the *l_o_* phase in the simple phase-separating GUVs. In exploring the basis of this drastic difference in lipid packing of the two leaflets, we used GPMVs, which have a more symmetric membrane^38^. Here the GP values before and after BSA-extraction are identical, settling at an intermediate value, at 0.49±0.07 (**Fig 3d**). These results strongly support the idea that the magnitude of difference of the GP of C3L, GP_T_(before) and GP_i_ (after back-extraction), is likely to be due to the compositional asymmetry of the two leaflets of the living cell membrane. These findings therefore reveal a highly ordered outer leaflet and a significantly more disordered inner leaflet, as also indicated from the earlier Di4-based microinjection studies of Lorent et al^14^.

To understand the basis of this extreme asymmetry, we altered the lipid compositions of the two leaflets by perturbing two lipid species, PS and SM, that are asymmetrically distributed. Depletion of inner leaflet localized PS causes both the leaflets to become more disordered^50^(**Fig 4c, d**). Given the transbilayer coupling propensity of the PS species and its high concentration in the inner leaflet ^50^, it is not surprising that loss of PS influences both the GP of the inner and outer leaflet. The effects of outer-leaflet SM depletion however remain restricted to the outer leaflet (**Fig 4a, b**). This was surprising considering that SM, with high melting temperature and long acyl chains would be predisposed to promote ordered/gel like domains and transbilayer coupling. It is likely that both the extent of depletion and the composition of the species that is depleted may play a role in the difference of these effects.

The transition of MDCK cells from a mesenchymal to an epithelial state (**M->E; Fig 5a, c**), presents a physiological example where compositional changes may be disproportionally reflected in membrane leaflets. Epithelial cells acquire increasing amounts of long chain, hydroxylated and saturated sphingolipid species as ascertained by mass spectrometry^28^. Consistent with the notion that composition modulates lipid packing during M->E transition we see an increase in the GP_T_ (**Fig 5e**). Importantly, this change is manifest in the outer leaflet since inner leaflet GP_i_ values are maintained unchanged during M->E. It is therefore tempting to speculate that the main protective barrier of the epithelial sheet, especially the apical domains, is formed by the tightly packed outer-leaflet while the more disordered inner leaflet provides sufficient flexibility to facilitate membrane protein function.

Focal adhesions (FAs) are large protein scaffolds responsible for cell-substrate adhesion and building the cellular machinery responsible for cell migration ^44^, and have been shown to have high GP^51^. This could be due to the recruitment of specific lipids to these sites or creation of a hydrophobic environment by the condensing of proteins at this site. Since C3L is able to directly report on lipid packing at each leaflet, our measurements at individual leaflet resolution shows that it is significant increases in GP_i_ that contributes to the higher order (GP_T_) that is characteristic of the FA site^51,52^. At the FA, integrin-signalling triggers the generation of hotspots of GPI-anchored protein nanoclusters^5^. These nanoclusters are formed due to immobilization of PS at the inner-leaflet by interaction with a dynamic actin-cytoskeleton, promoting transbilayer coupling with long-acyl chain containing GPI-anchored proteins^50^. This coupling should result in a more ordered local environment when templated from either leaflet^50^. We tested this hypothesis by deliberately templating the outer leaflet with defined patterns of crosslinking proteins created by immobilizing polyvalent antibody directed against a specific GPI-anchored protein. Consistent with the hypothesis we observe a larger increase in GP_i_ than observed for GP_T_ in regions that coincide with the patterned proteins (**Fig 6e-g**).

Studies on the diffusion of proteins tagged to lipids that partition into ordered/disordered membranes have indicated that antigen stimulation of the IgE-receptor (FcεRI) generates local order at the inner leaflet^53^. The reporter and the assay developed here provides a direct way to investigate these ideas, providing evidence for the same albeit in a different context. Similar mechanisms may function in the context of several signalling receptors, such as B and T-Cell receptor, and Integrin receptors, where receptor clustering is followed by signalling, accompanied by the formation of ordered membranes^11,42,51,53^. Such coupling may be a general property of self-assembling or self-organising transmembrane proteins with functional consequences of building distinct local leaflet-specific membrane domains due to their preference for specific protein and lipid species. The results presented here thus indicate that global membrane environment is set up by the specific composition of each leaflet resulting in a large difference between the membrane order between the outer and inner leaflet. It is likely that locally assembled protein scaffolds exploit this to create spatially confined transbilayer coupled domains, resulting in ordered domains which are correlated across the two leaflets. Furthermore, the design principles used for the chemical tools developed here are likely to have a more general application in leaflet-resolved imaging. This will open up new areas of investigations into biophysical principles that accompany construction of these membrane systems, and their biological function.

## METHODS

### Cell culture

Cells were maintained in their recommended media supplemented with NaHCO_3_ (Qualigens, Q14015), 10% heat inactivated fetal bovine serum (Gibco, 16000044) and non-essential amino acids along with an antimycotic-antibacterial cocktail (Sigma Aldrich, G1146) in sterile incubators at 37°C with 5% CO2 supply. CHOK1, IA2.2 (CHO cells over-expressing human folate receptor (Sharma et al, 2004)^54^, cells PSA3 cells (Kuge et al, 1986)^40^ were maintained in Ham’s F12 medium (AT144, Himedia). Mouse embryonic fibroblasts (MEFs) and MDCK cells were maintained in DMEM (AT170A, Himedia). All cell lines were passaged when the cells reached 70% confluency up to a maximum of 12 times. CHOK1 cells were used for the experiments unless otherwise mentioned.

### Spectra

Emission spectra of the compounds dissolved in the indicated solvents at 1uM concentration were collected at 5 nm steps while excited at indicated wavelengths on a Spectramax M5 Microplate reader, Molecular Devices.

### Giant Plasma Membrane Vesicle (GPMV) preparation

GPMVs were generated from 70% confluent CHO cells plated 48 hours prior to the experiment. Vesiculation was initiated by treatment with Paraformaldehyde and Dithiothreitol (PFA+DTT) as vesiculation agents using a protocol adapted from Sezgin et al, (2012). Briefly, cells were first labelled with C3L (10μM) for 5 mins at 37°C in M1G buffer (150mM NaCl, 5mM KCl, 1mM CaCl_2_, 1mM MgCl_2_, 20mM HEPES, 1mM Glucose, pH 7.4) under sterile conditions and washed 3x with M1G followed by 2x with GPMV buffer (10mM HEPES, 50mM NaCl, 2mM CaCl_2_, pH 7.4) under sterile conditions. The cells were then incubated in 1ml GPMV buffer containing vesiculation agents, 25mM PFA (15710, EMS) and 2mM DTT (D5545, Sigma Aldrich) for 1 hour at 37°C. The supernatant was decanted into an Eppendorf tube after giving a swirl to the flask to remove any attached GPMVs on the cell surface without detaching the cells. The GPMVs were allowed to settle for 20 mins at room temperature (RT). All imaging was done on the same day either directly or after doping with Fast DiI (D7756, Invitrogen) at 1:1000 dilution in GPMV solution from a 5mg/ml stock solution in ethanol. They were imaged inside a coverslip sandwich assembled by using a multi-layered double-sided sticky tape with punched holes to hold 20µl of GPMV containing solution on a 0.1% BSA (A3294, Sigma Aldrich) coated surface. The measurements for phase-separation and T_m_ in GPMVs at temperatures below 25°C was done on a temperature-controlled stage (model 9101 Polyscience) using a 40X air objective (0.75NA, Planfluor, Nikon Japan). Partition Coefficients were calculated by quantifying intensity of C3L and Fast DiI from the liquid disordered and ordered domains indicated by the preference of Fast DiI for liquid disordered regions.

### Giant Unilamellar Vesicle (GUVs) preparation

GUVs were prepared using the agarose (MB002, Himedia) gel swelling method (Horger et al, 2009)^30^. In brief, 1% agarose prepared in MilliQ water was put on a clean coverslip activated in an UV chamber supplied with oxygen flow just before adding agarose for 15 mins. The agarose coated coverslips were allowed to air dry in a hot air oven at 60°C for 30 mins. Lipid mix consisting of either 1:1:1 DOPC:DPPC:Cholesterol (850375P Avanti Polar lipids, 850355C2 Avanti Polar lipids, 47127-U Sigma Aldrich) or only DOPC in chloroform (G12305, Qualigen) was spread at total lipid concentration of 4mM on the dried agarose coated coverslips using the length of a Hamiltonian syringe. These coverslips were allowed to vacuum dry for 2 h before swelling the gel on the coverslip at 45°C using pre warmed MilliQ water at 7.4 pH for 15 mins. The solution containing the newly formed GUVs was then collected in an eppendorf, allowed to settle for at least 20 mins at RT before adding C3L to a final concentration of 1uM and/or Fast DiI diluted at 1:1000 in the GUV solution from a stock of 5mg/ml in ethanol. The GUVs were imaged on a temperature-controlled stage (INUG2H-TIZSH, Tokai Hit) at mentioned temperatures on coverslips coated with 0.1% BSA 20 mins prior to imaging.

### C3L cell labelling

Cells were plated on coverslip bottom dishes 48 h prior to the experiment conducted at 70% confluency. Cells were washed 2x with M1G. Samples were labelled for 5mins at 37°C with 1uM (unless otherwise mentioned) C3L, stock at 20 mM was maintained in anhydrous DMSO (D12345) and diluted in M1G in case of cells or other aqueous solvents as used for other sample preparations. Post labelling cells were given 3x washes with M1G and imaged in M1G at room temperature unless otherwise mentioned.

### BSA based Back Extraction

Samples labelled with a dye were imaged in the suitable aqueous buffer to acquire pre-extraction images. The buffer in these samples was then replaced with 1% defatted BSA (A8806, Sigma Aldrich) dissolved in the same buffer. No further washes or buffer exchanges were performed. The images for post-extraction condition were acquired either every 1min for the kinetics measurements or at the end of 15mins for steady state measurements.

### Lipid perturbations

1. Phosphatidylserine (PS) – CHO cells mutant for the PSA3 gene (PSA3 cells^40^) were grown for 72hours in media supplemented with or without ethanolamine (411000, Sigma-Aldrich) in media with dialysed serum. The cells grown without ethanolamine were depleted of PS as assessed by the reduction of annexin-staining in cells treated with Ionomycin according to Raghupathy et al^50^.
2. Sphingomyelin (SM) – Fumonisin B1 (32936, Sigma) was used to deplete Sphingomyelin from cells^39^. It was added to the media of the cells after allowing them to adhere to the coverslip for 3hrs post plating. Fumonisin B1 treatment was maintained alongside cells in media with no inhibitor as a control to deplete SM as determined earlier^7^.
3. 7-keto Cholesterol (7KC) treatment was performed adapting a published protocol(Rentero C. et al, 2008)^42^. Briefly, 7KC was supplemented into cells in complex with the vehicle MβCD at 0.38 mM. A 50mM 7KC (700015P, Avanti polar lipids) stock was prepared in ethanol. 2µl out of 10µl of this stock was added every 5 mins in 0.38mM 990µl mβCD (C4555, Sigma Aldrich) prepared in M1G and maintained at 90°C over 30mins. The mix was cooled and added to cells for 30 mins at 37°C.

### Focal adhesion imaging

Cells were transfected with mCherry-Paxillin using jet prime transfection reagent using manufacturer’s protocol 24h after plating to mark focal adhesions^5^. These cells were co-imaged with C3L along with the Paxillin labelled focal adhesions, both before and after BSA addition.

### Crosslinking of receptors on cell surface

Coverslips with pre-fabricated patterns of activated surfaces were coated with either Cholera Toxin B - AF647 or secondary antibody labelled with AF647(1:100) for 2hrs at 4°C followed by 7-8 times PBS washes. Subsequently the primary antibody (Mov18, 5ug/ml) against human folate receptor was incubated for 2hrs on these coverslips at 4°C followed by 7-8 times PBS washes. Two day plated cells were serum starved for 1h and trypsinised before adding them to coverslips (Cytoochips starter’s A). About 30000 cells were added and allowed to spread in serum containing media on the coverslips for 3.5hrs (IA2 cells) and 1h (MEFs) before washing the media out and labelling with C3L in M1G. These samples were imaged for C3L and AF647 tagged 2°ab/CTxB signal to record the pattern and degree of protein binding.

### Imaging

Live confocal imaging was done on a CSU W1 SORA (or where indicated a CSU22 mounted on Nikon TiE microscope with Andor iXon 897 cameras) spinning disk mounted on a Nikon Eclipse Ti2 microscope equipped with two Andor iXon 888 cameras. C3L was excited using a 405nm Vortan laser passing through a quarter wave plate ensuring circularly polarized excitation. Images for C3L were acquired on two channels simultaneously in the two cameras with the emission separated by a 470LP dichroic. The emission filters used for collection were 450/50nm(Ch1) and 500/40nm(Ch2). Images were acquired using Micromanager. A Back ground was acquired using imaging buffer. G Factor (GF) was acquired using C3L dissolved in DMSO. Images were acquired using a Plan Apo VC 100x 1.4 NA objective, or a Plan fluor 40X 0.75NA or a Plan Apo VC 60X 1.4NA (Nikon Japan).

### Quantification of GP

Images in the two channels corresponding to Ch1 and Ch2 for C3L were aligned and necessary background and GF corrections were applied. Image calculation for GP and GF was done using the following equations^31^:

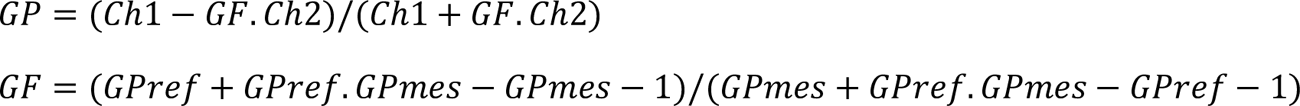

where,

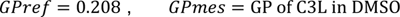

The derived 1024x1024 pixel images were cropped to quantify from the section of the frame presenting flat illumination by generating z project of all the generated GP colour maps and noting the section of the frame where the resulting LUT was white indicating independence of GP with respect to pixel position. Whole cell or 20x20 pixel ROIs were drawn on the cropped images while excluding very bright pixels in order to quantify GP values. Bright pixels were avoided because they could be structures like ruffles, endocytic pits or endosomes GP. Analysis was done using Fiji/ImageJ and Matlab.

### Derivation of the GP histograms

GPi = GP of inner leaflet GPo = GP outer leaflet

GP_T_ = GP of the membrane; weighted average GP of the two leaflets

Hence,

GP_T_ = α.GPi + (1-α)GPo ; α = fraction of dye at the inner leaflet

To derive the histograms in Fig. 5 and 6, 60 by 60 pixels ROIs were taken from the aligned images. GP_T_ and GPi images that were acquired via imaging before and after back extraction respectively. They were segmented using intensity based thresholding to remove the bright punctae or structures. The division of the corresponding total intensity (Sum of intensity acquired across Ch1 and Ch2) images before and after extraction yields α. Using an image based calculation and the derived parameters, GP_o_ images and values were derived. Using the histogram analysis function of ImageJ the respective histograms were then derived from the images

### Fluorescence correlation spectroscopy (FCS)

Fluorescence correlation spectroscopy (FCS) measurements were performed on a Zeiss LSM 780 equipped with a Confocor 3 System. Six sequential 10 seconds traces were obtained at the basal membrane of each of the cell using 6 µWatt from a 405 nm laser and a 40X 1.2 NA UV-VIS-IR C Achromat water-immersion objective. The correct focal distance was determined each time as the z-distance where the initial estimate of per particle brightness was highest. Next, the emission photon stream was recorded with the same objective, separated from the excitation, wavelength filtered between 411-553 nm to detect the whole emission range of C3L and detected on a gallium arsenide detector array. The intensity in time trace (*I(t)*) was autocorrelated into an autocorrelation curve G(τ) using the Zeiss onboard autocorrelator which calculates the self-similarity through:

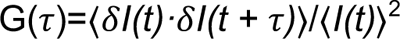

Here ‹› denotes the time-average, δ*I(t)* = *I(t)* - ‹δ*I(t)*› and τ is called the timelag. The raw autocorrelation functions (ACF) were fitted using a model with two timescales:

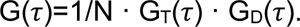

N reflects the number of moving particles in the confocal volume and G_T_(τ) is the correlation function associated to blinking/triplet kinetics:

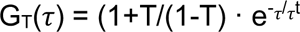

Where T is the fraction of molecules in the dark state (0.4±0.3) and τt corresponds to the lifetime of the dark state (1-200 µs). G_D_(τ) is correlation function associated to diffusion:

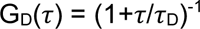

The relatively high non-correlated background lowers the signal to noise ratio as measured by counts per second per C3L. While increasing the excitation power will augment the signal to noise ratio there is an upper limit to increasing the power. Above excitation conditions of 10 µWatt, photobleaching of C3L starts dominating the diffusion time scales. At the used excitation condition of 6 µWatt there is a measurable signal from C3L. Nevertheless, the brightness per molecule (330±76 counts·s^-1^·molecule^-1^) make quantitative measurements challenging.

## Acknowledgements

We thank Andrey Klyemchenko (CNRS, Université de Strasbourg, France) for NR12S, Wolfgang Goldmann (Friedrich-Alexander-Universität Erlangen-Nürnberg, Erlangen, Germany) for primary MEFs and Y. Akamatsu (National Institute of Health, Shinagawa-Ku, Japan) for PSA3 cells. We thank all the members of the Mayor laboratory, in particular Sowmya Jahnavi and Sarayu Beri for suggestions and feedback on this manuscript. We also thank Lea Denker from Darius Koester’s Laboratory (University of Warwick) for helping with the GUV preparation protocol. We thank the NCBS Central Imaging and Flow Facility and Greeshma Pradeep S from Mayor lab for help with optical setups. C.P acknowledges doctoral fellowship support from National Centre for Biological Sciences, Tata Institute for fundamental Research. TSvZ acknowledges an EMBO fellowship (ALTF 1519-2013), a NCBS Campus fellowship and support from the Fundación General CSIC’s ComFuturo programme (European Union Horizon 2020 /Marie Skłodowska-Curie grant agreement No.101034263). M.S. and M.S. acknowledge support from CCRAS project: 1326/2022-23 (GAP3143). RAV and PPS acknowledge the support from the Council of Scientific & Industrial Research, India under Mission and MLP schemes (Project No. MLP5005). S.M. acknowledges support from Department of Biotechnology – Wellcome Trust India Alliance Margadarshi Fellowship IA/M/15/1/502018 and the Department of Atomic Energy, India (under Project No. RTI 4006), JC Bose National Fellowship (Department of Science and Technology, India), and Leverhulme International Professorship.

## Author Contributions

C.P. and S.M. designed the study and the probes. M.S, M.S and P.P.S contributed the synthetic strategy of the probes. The synthetic strategy was led by R.A.V and P.P.S. C.P. performed most experiments and analysed the data. T.S.v.Z performed FCS experiments and analysed the data. C.P and S.M wrote the paper with inputs and comments from all authors.

