## Supplementary figures and Table for "Solvatochromic reporter to image plasma membrane order leaflet by leaflet reveals a highly asymmetric bilayer locally modulated by transbilayer interactions"

**Suppl Fig. 1**

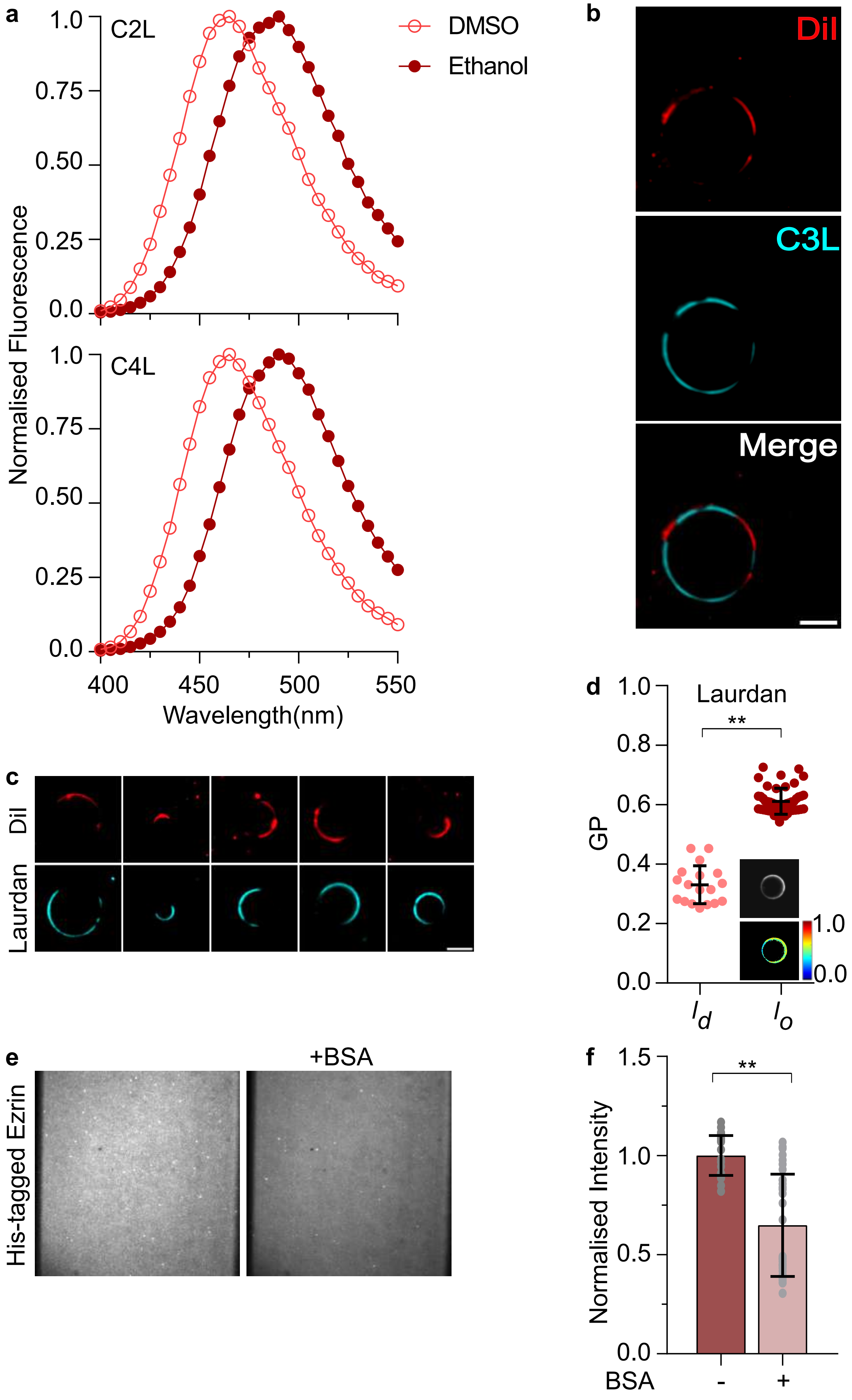

a
Normalised Fluorescence
C2L
C4L
DMSO
Ethanol
1.0
0.75
0.50
0.25
0.0
400
450
500
550
Wavelength(nm)
b
DiI
C3L
Merge
c
DiI
Laurdan
d
Laurdan
GP
1.0
0.8
0.6
0.4
0.2
0.0
**
$l_d$
$l_o$
1.0
0.0
e
His-tagged Ezrin
+BSA
f
Normalised Intensity
1.5
1.0
0.5
0.0
**
BSA
-
+

**Suppl Fig.S1**

(a)Emission spectra of C2L (top) and C4L (bottom) in DMSO and Ethanol at $\lambda$(ex)= 370nm. (b) Confocal images of GUVs showing fluorescence emission of Fast DiI (top), C3L (middle) and their merge (bottom) from the same vesicle at 25°C, prepared from 1:1:1 DOPC:DPPC:Chol lipid mix. (c) Confocal images of GUVs showing fluorescence emission of Fast DiI (top) and Laurdan (bottom) from the same vesicles prepared from 1:1:1 DOPC:DPPC:Chol lipid mix, at 25°C. (d) Graph showing GP values derived from Laurdan-labelled DOPC:DPPC:Chol vesicles determined from the Laurdan enriched ($l_o$) and depleted ($l_d$) regions at 25°C from atleast 12 vesicles. Inset shows confocal Laurdan emission intensity image(top) and GP image (bottom) of a 1:1:1 DOPC:DPPC:Chol vesicle. (e) TIRF images of 98:2 DOPC:[DGS-NTA(Ni2+)] bilayers doped with His-647SNAP-Ezrin protein before and after BSA-back extraction. (f) Graph showing distribution of His-647SNAP-Ezrin in the bilayer before (-) and after (+) back extraction with BSA. All data is plotted as mean±s.d. p values determined from unpaired t test Mann-Whitney test where ** indicates significance indicated by $p<0.005$. Scale=10µm.

**Suppl Fig. 2**

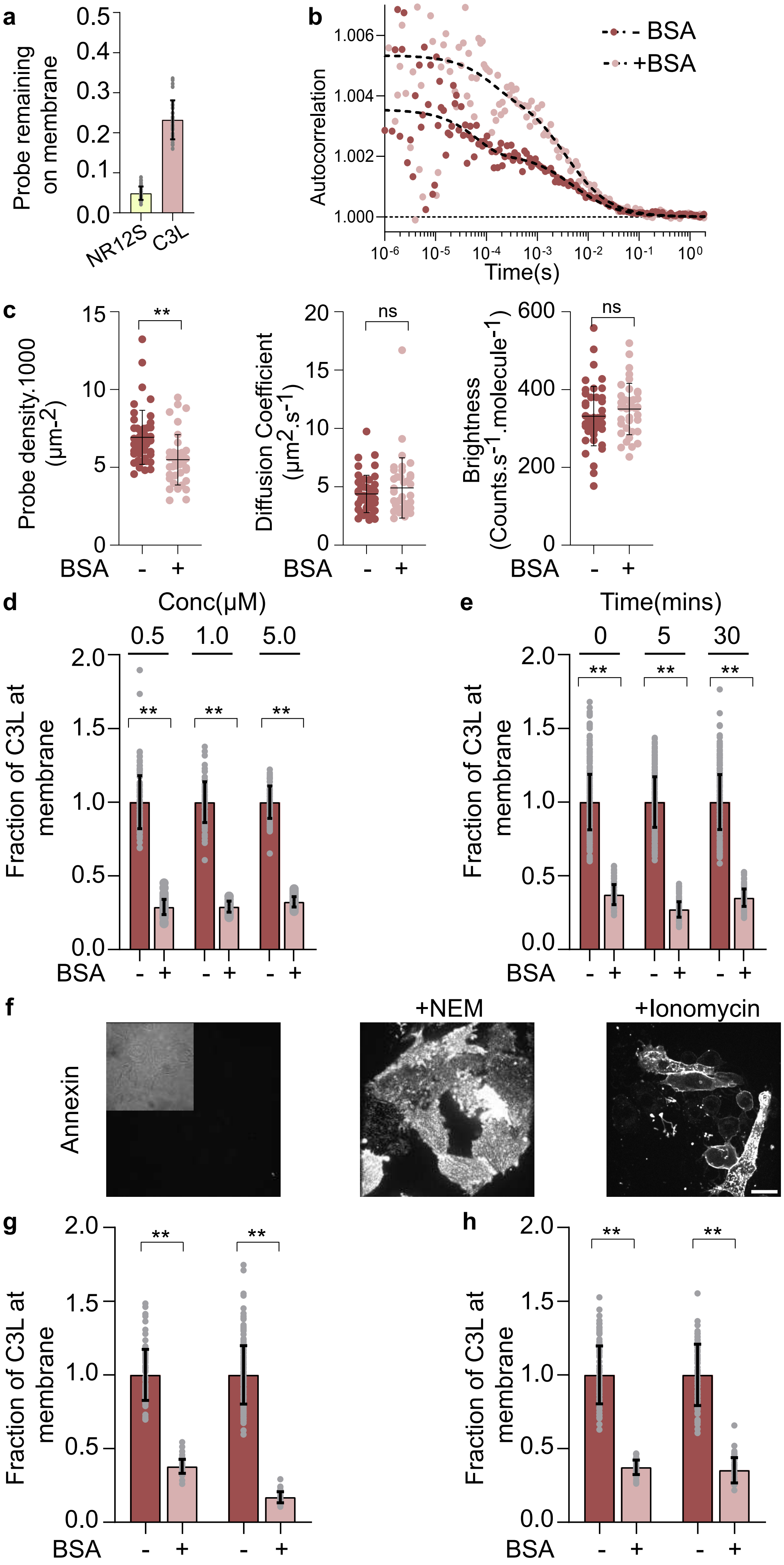

a
Probe remaining on membrane
0.5
0.4
0.3
0.2
0.1
0.0
NR12S
C3L
b
Autocorrelation
1.006
1.004
1.002
1.000
- BSA
+BSA
Time(s)
c
Probe density.1000 (µm-2)
15
10
5
0
**
BSA
-
+
Diffusion Coefficient (µm2.s-1)
20
15
10
5
0
ns
Brightness (Counts.s-1.molecule-1)
600
400
200
0
ns
d
Conc(µM)
0.5
1.0
5.0
Fraction of C3L at membrane
2.0
1.5
1.0
0.5
0.0
**
e
Time(mins)
0
5
30
f
+NEM
+Ionomycin
Annexin
g
h

**Suppl Fig.S2**

(a) Graph shows the fraction of probe (NR12S vs C3L) remaining at the cell surface (quantified at the rim) after back extraction performed as per the schematic in Fig2b. (b) Scatter plots of FCS auto-correlation with their corresponding fits (-----) before (-) and after (+) BSA-back extraction of C3L-labelled cells. (c) Graph shows quantification of density (left), diffusion coefficient (middle), and brightness (right) of C3L in cell membranes before (-) and after (+) back extraction. (d,e) Graphs show extent of extraction of C3L determined from fluorescence intensity obtained from confocal images of cells labelled with increasing concentration of C3L (d; data taken from at least 35 cells), or incubated over increasing periods of time at 37°C (e; data taken from atleast 60 cells) with 1µM C3L. Data in each column is normalized to the mean intensity of the same treatment before back extraction. (f) Confocal images show fluorescent intensity images of Annexin binding to untreated cells (left panel), cells treated with NEM, or ionomycin as indicated. The treatment conditions were the same as in Fig. 2c and Fig. 2e. Inset shows brightfield images of cells from the same field in left panel indicating the lack of any anexin staining in untreated cells. (g,h) Graph shows extraction efficiency (normalized fluorescence intensity) of NEM treated cells (g) and Ionomycin treated cells (h) as detailed in panels Fig 2c, e. Data is normalized to the mean intensity of C3L at the cell surface before extraction in the same treatment condition. All data is plotted as mean±s.d . *p* values determined from unpaired t test where ns indicates no significant difference and ** indicates significance indicated by $p<0.005$. Scale=10µm.

**Suppl Fig. 3**

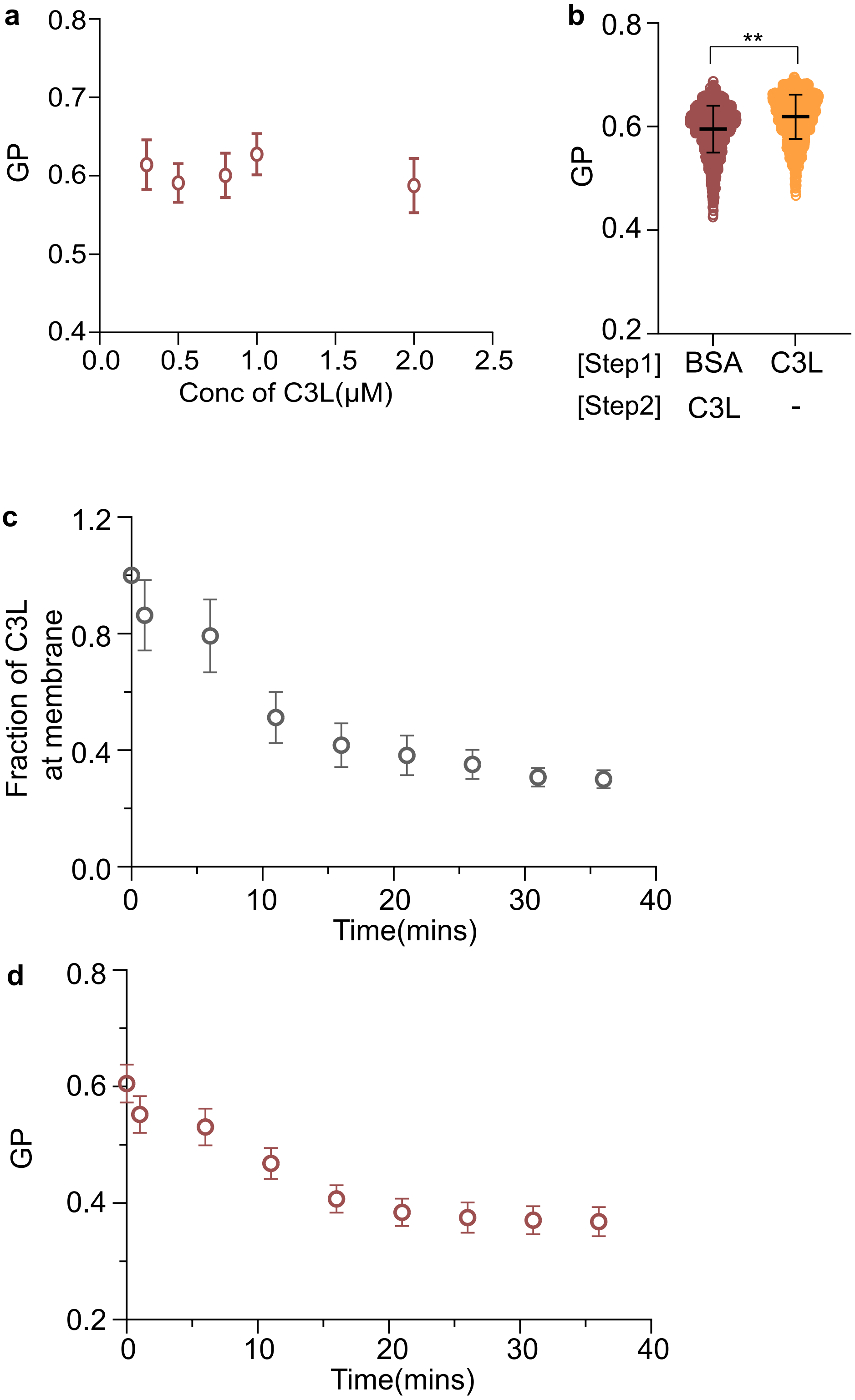
a
GP
0.8
0.7
0.6
0.5
0.4
0.0
0.5
1.0
1.5
2.0
2.5
Conc of C3L(μM)
b
GP
0.8
0.6
0.4
0.2
**
[Step1] BSA C3L
[Step2] C3L -
c
Fraction of C3L at membrane
1.2
0.8
0.4
0.0
0
10
20
30
40
Time(mins)
d
GP
0.8
0.6
0.4
0.2
0
10
20
30
40
Time(mins)

**Suppl Fig. S3**

(a) Graph shows GP values from cells labelled with indicated concentration of C3L from $\geq$ 45 cells. (b) Graph shows $GP_T$ values from cells pre-exposed to BSA based back extraction prior to labelling with C3L compared to cells directly labelled with C3L. Data taken from $\geq$ 100 cells. (c, d) Graph shows fluorescence emission intensity (c) and GP (d) of C3L at the cell surface as a function of time, from C3L-labelled cells exposed to BSA at t=0mins. The intensity data is normalized to that at t=0mins, and corrected for photobleaching. Each data point is obtained from 33 cells. All data is plotted as mean±s.d. *p* values determined from Mann-Whitney test where ** indicates significance indicated by $p<0.005$

**Suppl Fig. 4**

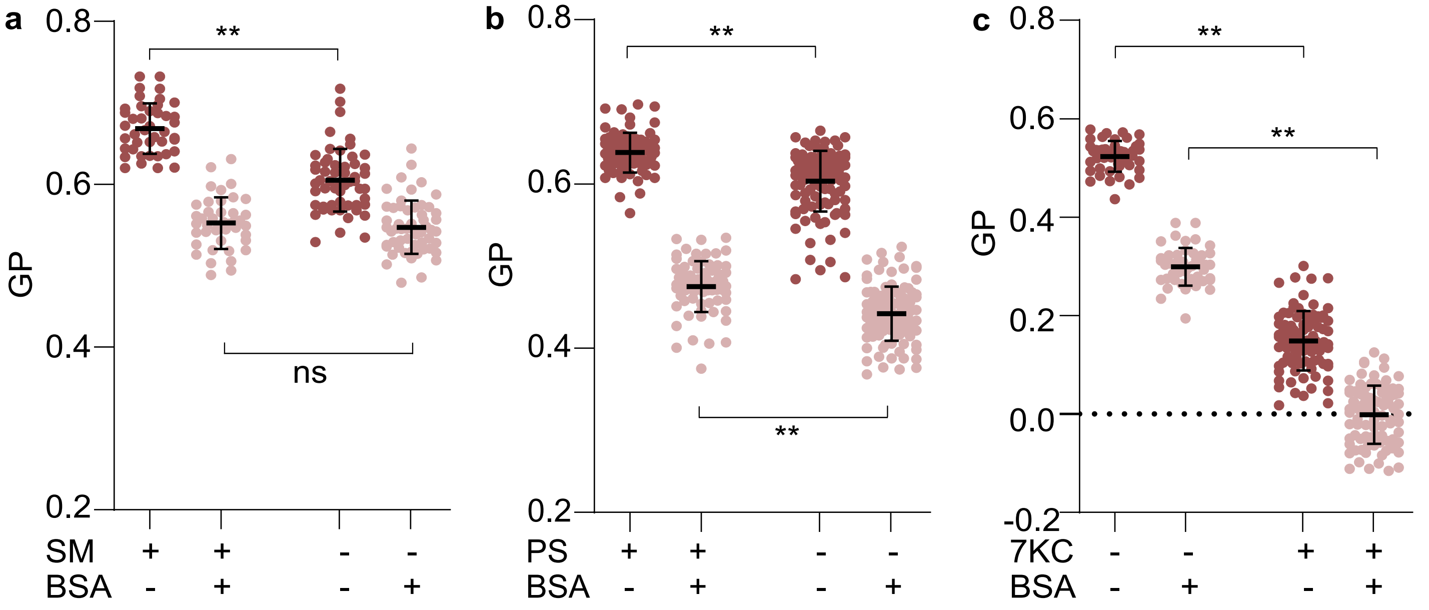


Suppl Fig. S4

(a-c) Graphs show the distribution of $GP_T$ (brown), and $GP_i$ (pink) from cells that were left untreated or grown with Fumonisin B1 to deplete Sphingomyelin from > 42 cells (a), from PSA3 cells maintained with and without ethanolamine to deplete PS levels from > 75 cells (b), or from cells treated with vehicle alone or along with 7-ketocholesterol to perturb the sterol distribution from > 55 cells (c). All data is plotted as mean±s.d, p values determined from unpaired t test where ns indicates no significant difference and ** indicates significance indicated by $p<0.005$

**Suppl Fig. 5**

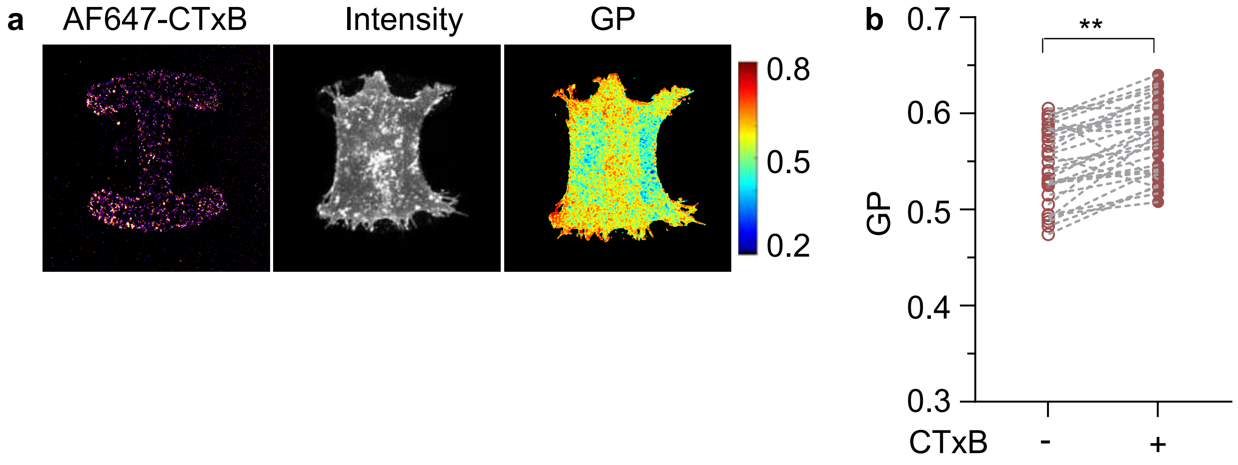


Suppl Fig. S5
(a) Confocal images of AF647 labelled Cholera Toxin B subunit (CTxB) covalently attached to patterned coverslips (left), C3L fluorescence emission intensity (middle) and GP maps (right) of cells plated on the coverslips (b) Graph showing GP values from the regions located on or off the cholera toxin patterns (+/- CTxB). Each datapoint is from a single cell with corresponding points on and outside the pattern of the same cell. Data is acquired from > 35 cells. All data is plotted as mean±s.d, p values determined from paired Wilcoxon non-parametric test where ** indicates significance indicated by p<0.005, *** indicates significance indicated by p<0.0001. Scale=10μm.

SUPPLEMENTARY MATERIAL

- **Table S1: Partition coefficient of C3L at indicated temperatures in GPMVs and GUVs**

| Temperature(°C) | | $K_p(l_o/l_d)$ |
|---|---|---|
| GPMV | 9 | 1.76 ± 0.3 |
| | 11 | 1.82 ± 0.18 |
| GUV | 25 | 3.36 ± 0.83 |

Data from Figure 1d, 1e.

Intensity based quantification of preference of C3L for $l_o$ and $l_d$ domains marked by the absence or enrichment of Fast DiI respectively.
